# Occipital Alpha-TMS causally modulates Temporal Order Judgements: Evidence for discrete temporal windows in vision

**DOI:** 10.1101/2020.03.30.015735

**Authors:** Samson Chota, Phillipe Marque, Rufin VanRullen

## Abstract

Recent advances in neuroscience have challenged the view of conscious visual perception as a continuous process. Behavioral performance, reaction times and some visual illusions all undergo periodic fluctuations that can be traced back to oscillatory activity in the brain. These findings have given rise to the idea of a discrete sampling mechanism in the visual system. In this study we seek to investigate the causal relationship between occipital alpha oscillations and Temporal Order Judgements using neural entrainment via rhythmic TMS. We find that certain phases of the entrained oscillation facilitate temporal order perception of two visual stimuli, whereas others hinder it. Our findings support the idea that the visual system periodically compresses information into discrete packages within which temporal order information is lost.

**Significance Statement:** Correlational evidence for a relationship between brain states and perception is abundant. However, in order to truly understand the causal relationship between the two, we need to be able to manipulate one and observe changes in the other. Neural entrainment via TMS serves as a valuable tool to interfere with cortical rhythms and observe changes in perception. Here, using rhythmic TMS-pulses at 10 Hz, we investigate the effect of the phase of entrained oscillations on performance in a temporal order judgement (TOJ) task. We observe that the causally entrained oscillation indeed modulates time perception rhythmically. On top of previous work on discrete perception we were able to 1. causally influence brain rhythms in a far more direct fashion using TMS, and 2. show that previous work on discrete perception cannot simply be explained by rhythmic fluctuations in visibility. In conclusion our findings support the long discussed idea that the temporal organization of visual processing is discrete rather than continuous, and is causally modulated by cortical rhythms. To our knowledge, this is the first study providing causal evidence via TMS for an endogenous periodic modulation of time perception.

## Introduction

Among neural oscillations, the occipital alpha rhythm is the most easily observed one (Berger, 1931). Since its discovery, it has been implicated in a vast number of cognitive functions such as attention, memory and inhibition (Bonnefond & Jensen, 2012; Foxe & Snyder, 2011; Jensen et al., 2012; Jensen & Mazaheri, 2010; Jokisch & Jensen, 2007; Klimesch, 1999, 2012; Klimesch et al., 2007; Tuladhar et al., 2007). A large body of literature investigates the effects of alpha amplitude on perception, linking high alpha power to high inhibition. More specifically, alpha power has been shown to increase in task irrelevant areas, whereas it decreases in task relevant areas, demonstrating its role in spatial attention (Foxe & Snyder, 2011; Kelly et al., 2006; Sauseng et al., 2005). Alpha oscillations play an even more dynamic role in the context of temporal attention, decreasing/increasing its amplitude at the moment when a target/distractor is expected (Rohenkohl & Nobre, 2011; van Diepen et al., 2015). On an even finer temporal scale we find that the phase of ongoing oscillations in the 5-15 Hz range influences perception. Busch et al. and Mathewson et al. (2009) demonstrated that the phase of occipital alpha oscillations is predictive of stimulus detection performance, implying that excitability in the visual cortex oscillates at around 10 Hz (Busch et al., 2009; Mathewson et al., 2009). These findings have been replicated several times using rhythmic entrainment at ~10 Hz via periodic visual stimuli or alpha-TMS (Dugué & VanRullen, 2017; Mathewson et al., 2010a; Romei et al., 2010; Spaak et al., 2014; Thut et al., 2011).

Many questions however remain. What is the exact relationship between rhythmic activity and perception? Are these periodicities mere changes in excitability, leading to fluctuations in detection probabilities or perceived intensities? Or do they impose discrete, non-overlapping windows, quantizing incoming visual information, akin to the snapshots of a camera? The former can be thought of as a “soft” version of periodic temporal perception whereas the latter can be thought of as a stricter “hard version” of periodic temporal perception.

The idea of a strictly discrete sampling mechanism (or “hard” version of discrete temporal perception (DTP)) states that the brain periodically divides the visual input into discrete windows or “perceptual moments” (Busch et al., 2009; Haegens et al., 2011; Lőrincz et al., 2009; Samaha & Postle, 2015; VanRullen & Koch, 2003; Vijayan & Kopell, 2012). Latterly the idea has gained support by psychophysiological and electrophysiological studies linking it to the occipital alpha rhythm (VanRullen, 2016). Recently it was shown that participant’s individual alpha peak frequencies are predictive of performance in a two-flash fusion task (Samaha & Postle, 2015). Following these lines it was demonstrated that visuo-auditory entrainment at the individual alpha frequency ± 2 Hz could facilitate or impair performance in a temporal segregation/integration task (Ronconi et al., 2018).

Further evidence comes from studies investigating the flash-lag effect (FLE), a visual illusion that has been suggested to arise from discrete sampling in the visual system (Chakravarthi & Vanrullen, 2012; Schneider, 2018). We demonstrated that visual entrainment at 10 Hz leads to a periodic modulation of temporal perception in the FLE (Chota & VanRullen, 2019). A key finding in the studies described above is that there is not only a fluctuation of perception (detection probabilities, perceived intensities), which is predicted by the soft version of DTP, but also a fluctuation of *time perception* itself (relative timing, temporal integration/segregation) which is the key prediction of the hard version of DTP. Given the findings described above we propose that the visual system discretely samples the visual scene at a ~10 Hz rhythm, analogous to the “hard” version of periodic perception (VanRullen, 2016). To test this hypothesis we utilized TMS to manipulate the occipital alpha rhythm and probed temporal order judgment (TOJ) at different phases of the entrained oscillation.

## Materials and Methods

### 1. Participants

25 participants (aged 18-31, 9 females) with normal or corrected to normal vision enrolled in the experiment. 7 participants had to be excluded during the first phosphene localization session because of their inability to see TMS-induced phosphenes, leaving 18 participants for the complete experiment and the final analysis. Note that this number of subjects excluded for this reason is very common in the TMS literature. Dropout rates of 40% due to inability to perceive phosphenes (at medial stimulation intensities) are commonly reported. Informed consent forms were signed before the experiment. The experiment was carried out in accordance with the protocol approved by the Centre National de la Recherché Scientifique ethical committee and followed the Code of Ethics of the World Medical Association (Declaration of Helsinki).

### 2. TMS apparatus, parameters and phosphene localization

The TMS stimulation was performed using a Magstim Rapid^2^ stimulator of 3.5 tesla, producing a biphasic current. At the beginning of the first session participants were tested on their ability to detect TMS-induced phosphenes. TMS stimulation was initiated at 55% of the maximum stimulator output applying 7 pulses at 20 Hz over occipital cortex. At the beginning of each phosphene localization trial participants were asked to fixate a central cross. Participants closed their eyes without changing the direction of their gaze. TMS was applied, participants opened their eyes and used the mouse to draw the outline and location of the perceived phosphene onto the screen. If no phosphene was perceived the coil position or stimulation intensity were changed manually and the procedure was repeated.

When a reliable phosphene was found the coil was fixated using an armed pedestal. We successfully elicited phosphenes in 18 out of 25 subject in the right (N = 9) or left (N = 9) visual field (Figure 1). Later, we used the phosphene location to place stimuli and scale them according to the cortical magnification factor (see section Stimuli below). Using a two-down one-up staircase procedure, we then determined the individual phosphene perception intensity threshold. Mean phosphene perception intensity threshold was 54.3% of maximum stimulator output. During the experimental TMS sessions we adjusted the TMS intensity to 75% of the individual phosphene perception intensity threshold (no subject reported perceiving a phosphene during the test trials).

**Figure 1.**
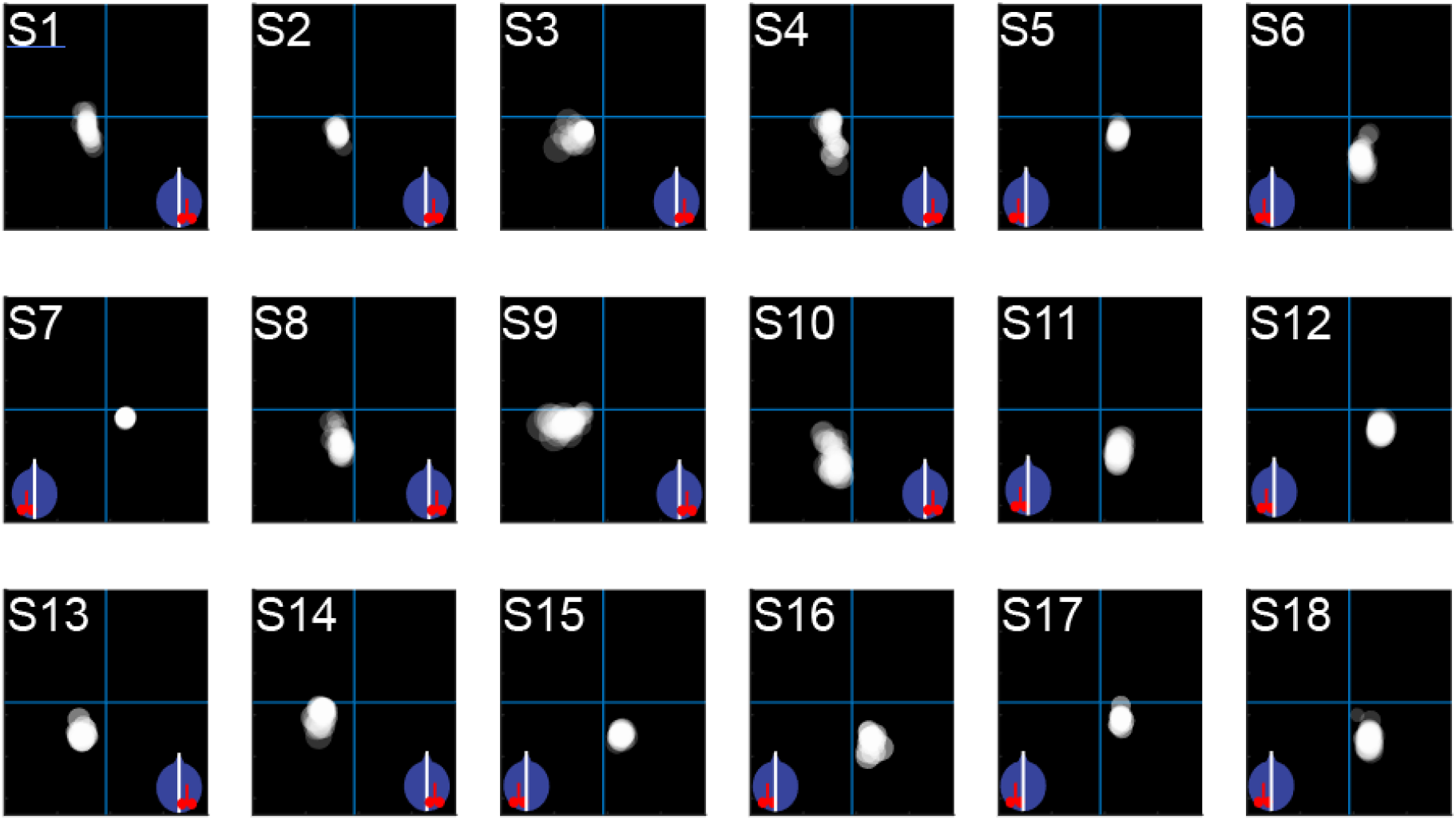
Stimulus locations for all subjects. The stimulus locations during the TMS blocks were determined during the phosphene localization. Participants had their eyes closed and received 7 TMS pulses at 20 Hz either over left or right occipital cortex. The red coil symbol over the blue head inset indicates the hemisphere of stimulation. The mouse was used by the subject to draw the outline of the perceived phosphene on the screen. Stimuli (here represented as superimposed white disks) were positioned inside the phosphene regions and scaled according to the cortical magnification factor (Horton & Hoyt, 1991). Note that due to slight changes in head position, the location of the phosphenes could change during the experiment. We re-localized phosphenes and adjusted the stimulus position at the beginning of every block.

### 3. Stimuli

Stimuli were presented at a distance of 57 cm with a LCD display (1920 x 1080 resolution, 120 Hz refresh rate) using the Psychophysics Toolbox (Brainard, 1997) running in MATLAB (MathWorks). Stimuli consisted of a central fixation cross (diameter = 0.3°), a square placeholder (black, 4°*4°), Gabor patches (Stim A and Stim B) of two orientations (45° and 135°, diameter = 3°, spatial frequency = 1.076 cycles/degree) as well as a mask in the form of a plaid (diameter = 3°), (Figure 2A). We included a masking stimulus in order to prevent subjects from using the persistence or the afterimage of Stimulus B to solve the TOJ task. Stimulus ***A*** will always be referring to the first stimulus of the sequence, whereas Stimulus ***B*** or "target" will be referring to the second stimulus of the sequence, irrespective of their orientation. During the task Stimulus ***A*** and ***B*** were presented in quick succession and participants gave Temporal Order Judgements by identifying the orientation of stimulus ***B***. Additionally we included a single stimulus condition where only one stimulus (Stimulus ***A*** or Stimulus ***B***) was presented. Participants reported these trials using a third button. The single stimulus condition was included in order to control for the possibility that TMS entrainment could lead to a significant decrease in stimulus visibility for one of the two stimuli (***A*** or ***B***). In that case we would expect an increase in single stimulus reports. We will show later in the behavioral results that this was not the case.

**Figure 2.**
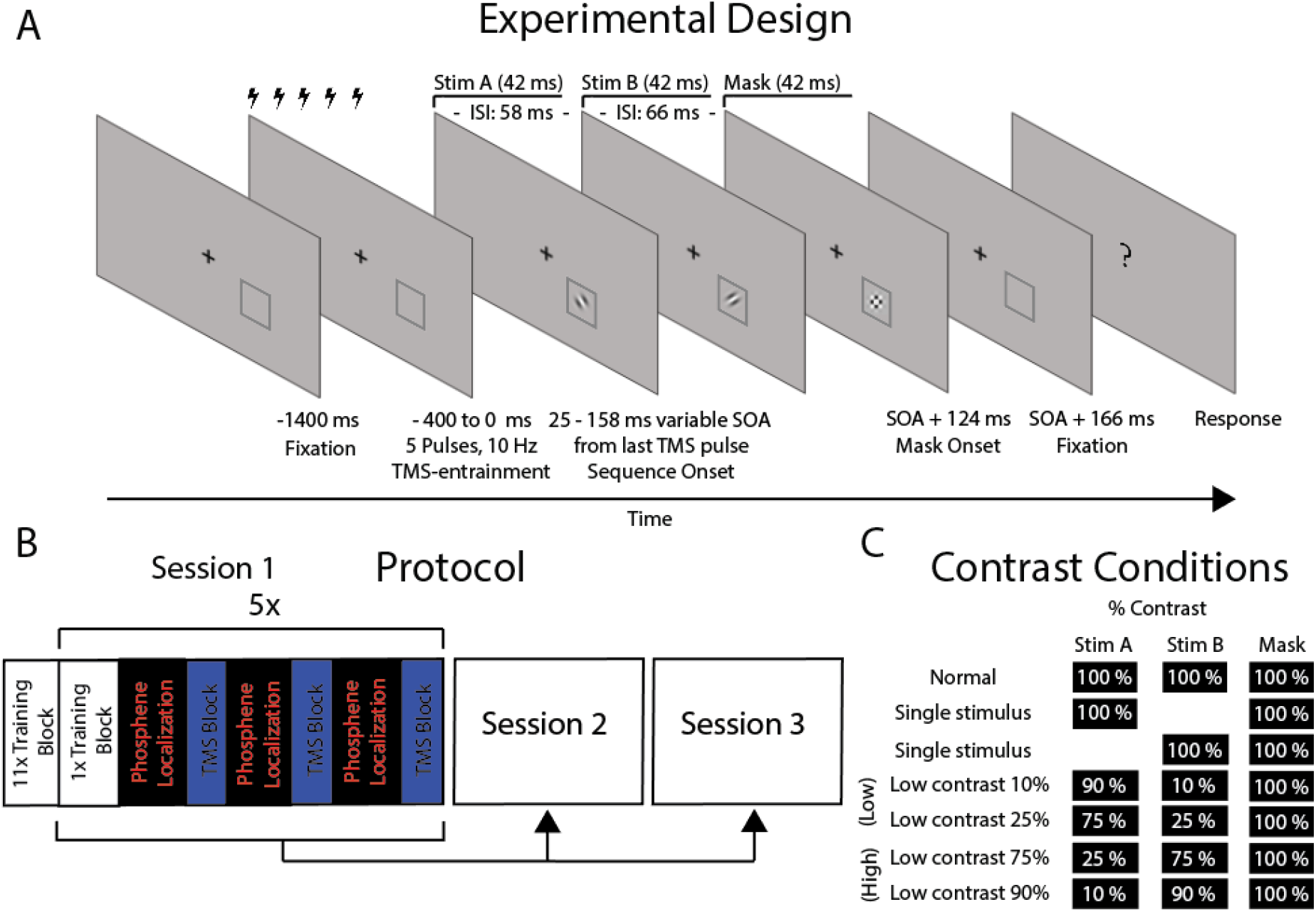
Paradigm. **A.** Experimental Design. Participant’s initiated trials via button press while fixating at a central fixation cross. After 1000 ms 5 TMS pulses at 10 Hz were administered. We probed TOJ performance at nine time points between 25 ms and 158 ms after the last TMS pulse. This was done by presenting a sequence of two Gabor patches with different orientation (Stim A and Stim B, 45° or 135°, stimulus length: 42 ms, ISI 58 ms), followed by a mask to prevent stimulus persistence or long afterimages. Participants had to identify the orientation of the second stimulus which was different on every trial. Alternatively participants could report (using a different key) a single stimulus condition where either the first or second stimulus was omitted (see C.). In the TMS Blocks no feedback was provided. **B.** Protocol. The experiment was performed on 3 separate days within 5 days. On the first session only, participants performed an extensive training consisting of 253 trials without TMS, with feedback and with a predefined stimulus location. Afterwards the main experiment was performed consisting of 5 repetitions of the Block sequence indicated in the figure. These 5 repetitions were repeated on session 2 and 3. **C.** In order to control for an effect of TMS on the visibility of the stimuli we used 5 different contrast conditions. In a subset of trials the contrast of the stimuli in the sequence was separately reduced to either 10%, 25%, 75% or 90%. The contrast of the Mask was identical in all conditions. Additionally we included a single stimulus condition where one stimulus was omitted.

The experiment consisted of a training condition and a TMS condition. We included a rather extensive training session in order to minimize any potential practice effects that could influence our TMS sessions and to reduce potential biases towards stimulus visibility, as we will explain later. The training condition was run with the parameters described above. The location of placeholder and stimulus sequence was fixed in the lower right or left visual field with equal probability (eccentricity 7°). The stimulus parameters in the TMS condition were identical to the training condition except for the size and position of the stimuli and placeholder which were adjusted based on the phosphene location acquired during the phosphene localization described in the previous paragraph. The location of the stimuli was chosen to be within the reported phosphene area, and as close to the position of the training stimuli as possible (Figure 1). This ensured that stimuli in the TMS condition were not presented further than 7° visual angle from the position of the training stimuli in 96% of trials to keep training and TMS conditions comparable. The stimuli were scaled based on their eccentricity according to cortical magnification (Horton & Hoyt, 1991) in order to match the cortical representation of the stimulus to the actual cortical stimulation site during TMS. For example, the stimulus diameter measured 3° at 7° eccentricity and 3.78° at 10° of eccentricity. The contralateral stimulus position was determined by flipping the ipsilateral position around the central y-axis. Left and right stimulus trials were matched pseudo-randomly. Stimuli were presented on a gray background.

We modulated the contrast of our stimuli in a subset of trials. This was done for two reasons. First of all, it allowed us to control for the potential effects of TMS on the visibility of the stimuli. We hypothesized that any detrimental effects of TMS on visibility should be maximal when target contrast is minimal. More specifically, this potential confound should manifest as a reduction in performance for stimuli that were presented on the hemisphere contralateral to TMS, with high contrast stimuli being least and low contrast stimuli being most affected. As we will show later in the Results section, this was not the case. Second, by having stimuli vary in contrast, we aimed to discourage participants from basing their TOJ judgments on visibility (e.g. always judging the most visible stimulus as occurring first or last), by making stimulus visibility inconsistent across trials while at the same time providing feedback. Therefore, even if the entrainment oscillatory phase should affect visibility, participants are actively encouraged to neglect visibility cues and base their decision purely on the perceived temporal order. For this second reason we decided to include several intermediate contrast conditions to better simulate perceptual ambiguity and increase the sense of unreliability of stimulus visibility. 5 Contrast conditions were used (Figure 2C): Normal (100% contrast for ***A*** and ***B***), low contrast 10% (Stimulus ***A***: 90% contrast, Stimulus ***B***: 10% contrast), low contrast 90% (Stimulus ***A***: 10% contrast, Stimulus ***B***: 90% contrast), low contrast 25% (Stimulus ***A***: 75% contrast, Stimulus ***B***: 25% contrast), low contrast 75% (Stimulus ***A***: 25% contrast, Stimulus ***B***: 75% contrast). Condition names (e.g. low contrast 75%) refer to the contrast value of the second stimulus of the sequence (Orientation of Stimulus B which had to be identified in the TOJ task) and will be used for later reference. Low percentage values represent low contrast or small difference to background luminance. The percentage values represent the Michelson contrast, defined as the ratio between the minimal/maximal luminance value of the stimulus and the gray background ((max. Luminance – min. Luminance)/(max. Luminance + min. Luminance)).

### 4. Protocol

On the first session only, participants performed an extensive pre-training consisting of 12 Training Blocks (23 trials each). Training blocks were identical to the TMS blocks with few exceptions. No TMS-pulses were applied during training and the placeholder as well as sequence presentation was in the lower right or left (equally balanced) visual field. Participants received positive or negative feedback in the form of the fixation cross turning green or red. One goal of the Training Blocks was to prevent participants from basing their TOJ judgements on the visibility of stimuli by making the contrast maximally uninformative as described above. Furthermore, it served to keep performance on a steady level by providing frequent feedback to the participant. In the training blocks 44.3% of trials were low contrast trials, 22.3% were single stimulus trials and 33.3% were normal contrast trials.

Afterwards subjects performed the main experimental procedure, which was repeated on session 2 and session 3. In the main experimental procedure participants performed 19 blocks (15 TMS blocks, 5 Training blocks) with 23 trials per block (In the last TMS block only 20 trials were collected). 3 TMS blocks were interleaved with 1 Training Block. Over the course of 3 sessions this resulted in 1035 trials in the TMS and 345 trials in the training condition per subject (50% trials contralateral to TMS, 50% trials ipsilateral to TMS). Per SOA and per visual field 57 trials were collected. Before each of the 45 TMS blocks participants performed a phosphene localization to make sure that the coil position had not changed and the correct cortical area was stimulated. Trials started with the central fixation cross and the placeholder on either side of the screen (Figure 2 A). The placeholder served as a cue to indicate location of the stimulus sequence with 100% cue validity. This served to avoid potential attentional confounds by always directing attention to the precise location of the upcoming stimuli. Participants initiated the trial via button press. 1000 ms after the button press 5 TMS pulses (100 ms between pulses) were administered over the course of 400 ms. Starting with the last TMS pulse, after a variable delay (25 – 158 ms in steps of 16.7 ms) the stimulus sequence was presented inside the square placeholder. The orientation of Stim A and Stim B was pseudo-randomly chosen every trial (45° and 135° or vice versa).The presentation length for Stim A, Stim B and the mask was 42 ms each. The ISI between Stim A and Stim B was 16 ms. The ISI between Stim B and the mask was slightly longer with 24 ms. During Piloting we observed a forward and backwards masking effect on the second stimulus which we compensated by shifting the onset of the mask to a later time-point. This was done to equalize visibility between Stim A and Stim B which we verified during piloting in 4 subjects. After a delay of 1000 ms participants reported either the orientation of Stimulus B (arrow key left or right) or reported perceiving a single stimulus (arrow key up). Trials were randomly chosen from the normal condition (66.3% probability) [Stim A (normal contrast), Stim B (normal contrast), Mask], low contrast condition (22.3% probability) [Stim A (low contrast), Stim B (low contrast), Mask] or a single stimulus condition (11.3% probability) [Stim A (normal contrast), omitted, Mask] or [omitted, Stim B (normal contrast), Mask] (See Figure 2C). During the phosphene localization and the experiment, the participants head was fixed between a chinrest and the TMS coil, leading to a stable position. In total, a maximum of 2215 TMS pulses were administered per session.

### 5. Data Analysis

We analyzed the data collected during the training blocks and the TMS-blocks separately. The 11 pre-training blocks at the beginning of session 1, which were included to prevent visibility biases, were not included in the analysis because of potential practice effects. Only training blocks that were collected during the main experimental procedure (interleaved with the TMS blocks) are included in this analysis (3 × 5 blocks of 23 trials per subject).

#### 5.1 Training Blocks mean performance

Due to the significantly smaller number of trials collected in the training blocks compared to the TMS-blocks we did not analyze the time course of TOJ performance during training but looked at the overall performance for all 6 conditions (*normal contrast* condition, 4 low contrast conditions, *single stimulus* condition). The 4 low contrast conditions (Figure 2C) were merged into two groups based on the contrast of the stimulus B (target), separating them into a *low target contrast* group (Stimulus B contrast 10% and 25%) and *high target contrast* group (Stimulus B contrast (75% and 90%). Note that the major aim of including several fine grained contrast conditions in the first place was to counteract potential visibility biases, by preventing subjects from relying on visibility instead of TOJ. The separation based on Stimulus B contrast was done to control for possible effects of TMS on Stimulus B (target) visibility which should be maximal for the low contrast stimuli (for which orientation was reported) in the *low target contrast* group. Mean performance (Hits/Misses) was averaged within each subject for 1. *normal contrast*, 2. *low target contrast* (low contrast 10%, low contrast 25%), 3. *high target contrast* (low contrast 75%, low contrast 90%) and 4. *single stimulus* conditions separately. For the low contrast conditions we acquired 41.9 trials (*low target contrast*) and 42.2 trials (*high target contrast*) on average per subject. For the *normal contrast* and the *single stimulus* conditions on average 81.3 and 29.5 trials were collected per subject. The behavioral performance for each condition during training was quantified using a repeated measures ANOVA.

#### 5.2 TMS Blocks

##### 5.2.1 Analysis of mean performance

The analysis of the mean performance for the 6 conditions (*normal contrast* condition, 4 low contrast conditions, *single stimulus* condition) was done similarly to the analysis in the previous section by merging low contrast conditions into 2 groups (*low target contrast*, *high target contrast*). Mean performance per subject in the resulting 4 conditions and for each visual field (contraTMS and ipsiTMS) was analyzed using a repeated measures ANOVA.

##### 5.2.2 Analysis of TOJ time-series

For our main results, the effect of TMS on subsequent TOJ-performance was analyzed as a function of the delay between the last TMS-pulse and presentation of the first stimulus of the sequence. Since the other contrast conditions only served to ensure that participants engaged in unbiased TOJ judgments only the normal contrast condition (66.3% of trials) was included in this analysis. We removed all responses where subjects erroneously reported single stimuli (5.4%), counting only left/right responses (making chance performance 50%) .We probed TOJ performance at 9 time points (SOA’s) after TMS (25 ms, 41.7 ms, 58.3 ms, 75 ms, 91.7 ms, 108.3 ms, 125 ms, 141.7 ms, 158.3 ms). On average 712.5 trials (24.2 SEM) were collected per subject. Performance (Hits/Total number of trials) was averaged within each subject at each of 9 SOA’s for stimuli presented contralateral and ipsilateral to TMS respectively. The resulting time series were normalized by subtracting the mean and averaged over subjects, resulting in a grand average contralateral (contraTMS) and ipsilateral (IpsiTMS) time series.

In an additional analysis the individual 133 ms long TOJ time-series were subtracted on a subject by subject basis in order to calculate the difference waves between the two conditions. These difference waves were averaged resulting in the grand average difference wave (contraTMS minus ipsiTMS). Time-series were zero-padded to a length of 6 times the original window length (6*133.3 ms) and analyzed in the frequency domain using FFT (frequency resolution 1.1 Hz). 9 SOA’s at 60 Hz allowed for a Nyquist frequency of 30 Hz. The complex FFT coefficients were squared to obtain oscillatory power at each frequency. To statistically test if the time-series contain significant oscillatory power we calculated 1.000.000 surrogates by shuffling the SOA-labels between trials for every subject, and repeating all analysis steps for each surrogate. The original power-spectrum was then compared to the surrogate distribution and p-values were corrected for multiple comparisons using the False Discovery Rate. The FFT revealed a peak at 10 Hz for contraTMS and a peak at 7.8 Hz for the ipsiTMS condition. For the phase analysis, we therefore decided to extract individual phase angles from the center frequency 8.9 Hz component of the FFT of the down-sampled and normalized contraTMS and ipsiTMS time-courses as well as the contraTMS minus ipsiTMS difference wave. Individual contraTMS and ipsiTMS phase angles were subtracted (contra minus ipsi, pairwise subtraction) to investigate the phase relationship in individuals. Rayleigh’s test for non-uniformity was used to statistically test if individual phases were significantly coherent.

## Results

In this study we seek to provide causal evidence for the hard theory of discrete perception. Specifically we aim to show that discrete perception entails a periodic compression of time information. Our hypothesis therefore states that within a perceptual moment, perception of the temporal order of events is impaired. To test this hypothesis we entrained participants’ alpha oscillations using 10 Hz-TMS over early visual areas. We probed TOJ performance, an index of perception of relative timing, at different phases of the entrained oscillation after stimulation. A significant oscillatory component at 10 Hz at the entrained location (as measured by a frequency analysis on the average TOJ time-course) would suggest a rhythmic modulation of time perception.

### 1. Mean performance and low contrast control conditions

Mean performance in the normal and low contrast conditions was compared between Training blocks (no-TMS) and TMS blocks. During Training the mean performance across subjects and visual fields was 63.14% (±3.5% SEM) for the *normal condition*, 56.4% (±4.4% SEM) for the *low target contrast* condition, 63.1% (±3.4% SEM) for the *high target contrast* condition and 49.9% (±6.6% SEM) for the *single stimulus* condition. In the TMS blocks the mean TOJ performance across observers in the *normal contrast* condition was 64.4% (± 2.6% SEM) for stimuli contralateral and 62.4% (± 2.5% SEM) for stimuli ipsilateral to the cortical entrainment site. For the *single stimulus* condition performance was 54.7% (± 5.8% SEM) for stimuli contralateral and 53.3% (±7.1% SEM) for stimuli ipsilateral to the cortical stimulation site. In the *low target contrast* condition average performance was 63.6% (±3.3% SEM) for the contraTMS condition and 57.4% (±3.7% SEM) for the ipsiTMS condition. Last, in the *high target contrast* condition performance was 65.8% (±3% SEM) for the contraTMS and 62.1% (±1.8% SEM) for the ipsiTMS condition.

The proportion of “single-stimulus” responses is an indicator of subjects’ ability to discern the presence of two stimuli, and therefore to perform genuine temporal order judgments rather than base their responses on the most visible stimulus. We calculated d’ by comparing the proportion of single-stimulus responses in the single-stimulus condition (hits: 51.9%) to the proportion of single-stimulus responses in the normal contrast condition (false positives: 5.4%). Sensitivity was d’ = 1.7 indicating that participants were well able to differentiate between 2 stimuli and 1 stimulus conditions. Furthermore, the single stimulus report rate of 5.4% in the normal contrast condition indicates that participants could perceive the 2 stimuli in the vast majority (94.6%) of normal contrast trials, and suggests that TMS-induced performance fluctuations stem from fluctuations in temporal order perception, rather than stimulus detection.

To reinforce this conclusion, both TMS blocks and Training blocks were analyzed using a repeated measures ANOVA with factors TMS (TMS versus Training), hemisphere (contra versus ipsilateral to TMS) and contrast (normal, low target contrast, high target contrast, single stimulus). We were particularly interested in verifying that any possible TMS-induced changes in TOJ performance were not mediated by a modulation of the visibility of the stimuli (and specifically, the target stimulus), as this could have been a confounding factor. We hypothesized that if TMS had a detrimental effect on stimulus visibility, this should be reflected in average performance differences when comparing TMS and Training (no-TMS) trials. Furthermore, these differences should be even more pronounced in low contrast conditions, where the target visibility is closer to the visibility threshold. The repeated measures ANOVA revealed a main effect of hemisphere (F(1,17) = 7.6, p = 0.013) but not of TMS (p = 0.38) or contrast (p = 0.1). Post hoc tests revealed significantly higher performance in the contralateral compared to the ipsilateral condition (p = 0.01), in both visual fields for TMS and Training conditions. The increased contralateral performance might be the result of a general attentional bias towards the hemisphere where the phosphene was located, irrespective of TMS stimulation. Most importantly we did not find an interaction effect between factors TMS and hemisphere (p = 0.55) or TMS, hemisphere and contrast (p = 0.73) speaking against the possibility that TMS might influence target visibility. Finally we also investigated the relative number of single stimulus responses for all conditions. We hypothesized that if TMS leads to an occlusion of a single stimulus this should be apparent in an increase in the ratio of single-stimulus responses. Moreover this should especially be the case for stimuli presented contralateral to TMS. The repeated measures ANOVA revealed a main effect of contrast (F(3,51) = 71.1, p < 0.005) but not of TMS (p = 0.12) or hemisphere (p = 0.69). Post hoc tests revealed significantly different ratios of single stimulus reports between all contrast conditions (Normal contrast: 5.4%, high target contrast: 10.9%, low target contrast: 28.8%, single stimulus condition: 51.9%, p&<0.05 respectively). This finding comes as no surprise and shows that our contrast manipulation successfully created distinct levels of target visibility. Importantly however no interaction effect between TMS and hemisphere (p = 0.2), TMS and contrast (p = 0.73) or TMS, contrast and hemisphere (p = 0.1) was found, indicating that TMS did not lead to a higher number of single stimulus responses. To avoid confusion, we would like to mention again that, although an increase in single stimulus responses should lead to a decrease in overall performance, the results of both ANOVA’s are not contradictory, since in the first ANOVA the single-stimulus reports were removed in all but the single stimulus condition.

We interpret the above presented findings as evidence that TMS did not have a general, time-unspecific effect on target visibility or detection, which could have indirectly led to a modulation of TOJ performance, but rather modulated perceived relative timing of the stimuli.

### 2. α-TMS leads to periodic modulation of TOJ performance

In order to verify that TOJ performance was rhythmically modulated by α-TMS we calculated the average TOJ time-series over individuals. This was done by calculating the average performance for each of the 9 SOA’s. Performance was analyzed separately for sequences presented contralateral (contraTMS) and ipsilateral (ipsiTMS) to α-TMS. Only normal contrast trials (66.3% of all trials) were included in this analysis, in order to discard any possible influence of contrast differences on stimulus visibility. For this analysis we excluded trials in which participants reported only a single stimulus: when both stimuli are visible, we can reasonably assume that subjects’ reports reflect their perceived temporal order, as per the task instructions. TOJ time-series were normalized by subtracting the individual mean before applying the FFT. Figure 3A shows the original, un-normalized time-series. Initial inspection of the time course for contraTMS indicates a strong oscillation in the TOJ performance lasting for more than 158 ms. To quantify this effect we performed a frequency analysis on the average TOJ-time -series. The resulting power spectrum revealed a dominant oscillation at 10 Hz (Figure 3D). For statistical validation of this peak we created 1.000.000 surrogates by shuffling the 9 SOA-bin labels within subjects and recalculated the power spectrum of the resulting TOJ time series. P-values were computed as the percentile of the mean power values within the bootstrapping distribution. This allowed us to test the null-hypothesis that the power spectrum of the average TOJ time-series does not show a peak at a specific frequency. The TOJ time-series power at 10 Hz was significantly higher compared to the surrogate distribution (Figure 3D, 10 Hz: p = 0.00006, FDR corrected). We analyzed the ipsiTMS TOJ time-series in an identical fashion. Initial inspection indicated a slightly weaker oscillation peaking at a lower frequency of 8 Hz (Figure 3B). The frequency analysis and statistical analysis of the ipsiTMS time-series showed significantly more power compared to the surrogate distribution (Figure 3E). Power peaked at 8 Hz (p = 0.0039, FDR corrected) but was still significant at the entrainment frequency of 10 Hz (p = 0.015, FDR corrected).

**Figure 3.**
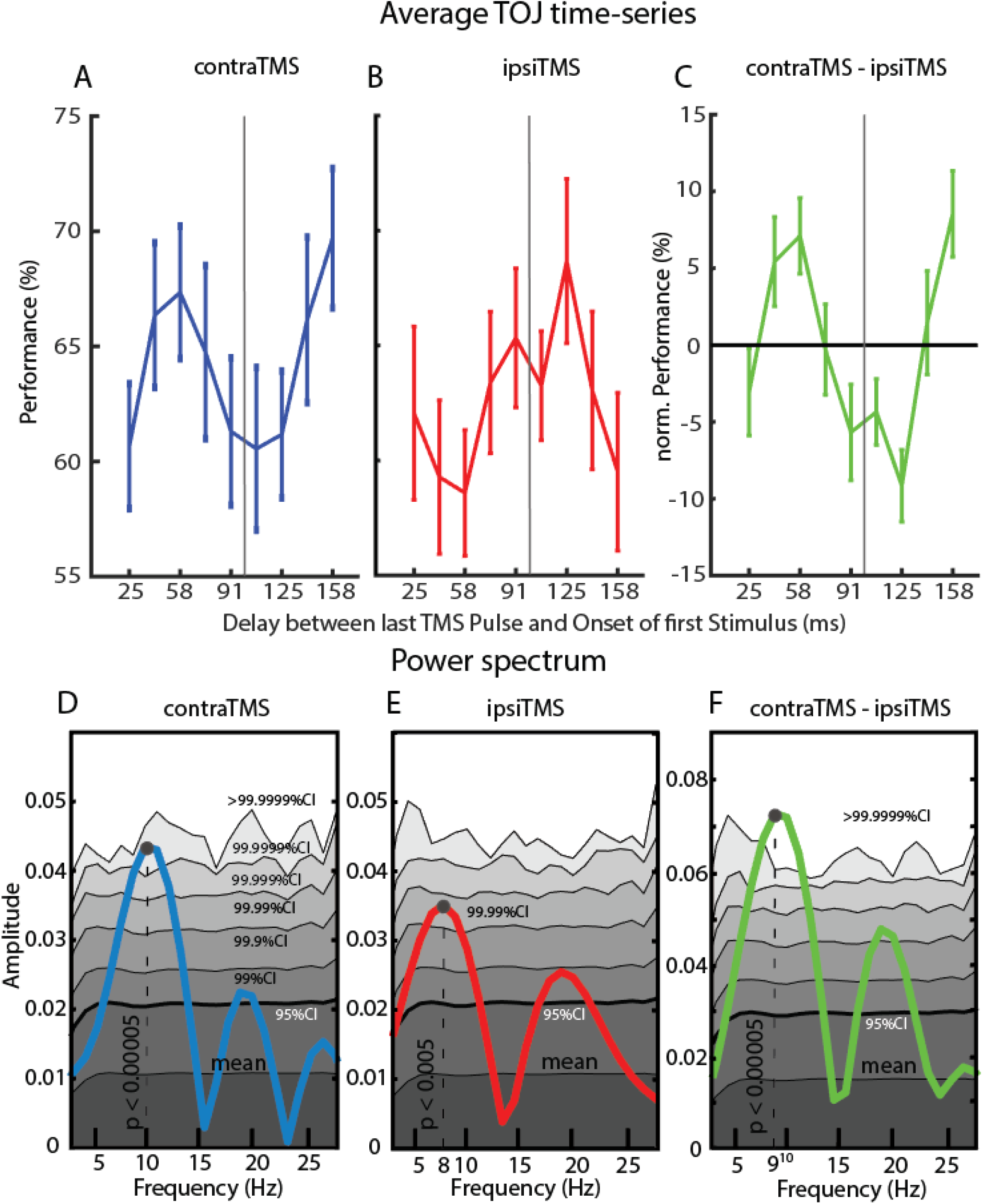
Main findings. **A.** Average TOJ time-series (N = 18) for stimuli presented contralateral to TMS. Error bars represent standard error of mean. The vertical gray line indicates the hypothetical time point of the next TMS pulse if the entraining sequence had continued. **B.** TOJ time-series for ipsilateral stimuli. **C.** Difference wave between contraTMS and ipsiTMS time series. **D.** Power spectrum of the contraTMS time-series. We compared the peak at 10 Hz to a surrogate distribution (1.000.000 surrogates) which revealed a significantly higher power at this frequency compared to other frequencies after correcting for multiple comparisons (p = 0.00006, FDR-corrected: p = 0.005). Colored areas: Dark gray: Mean of the surrogate distribution; Light gray: 95% Confidence Interval; 99% CI; 99.9% CI; 99.99% CI; 99.999% CI; 99.9999% CI; White: >99.9999% CI. **E.** Power spectrum of the ipsiTMS time-series. We observed a relatively strong oscillatory component at around 8 Hz. Analogous to (D) we analyzed oscillatory power at 8 Hz and found significantly more power compared to the surrogate distribution (p = 0.00038, FDR-corrected: p = 0.0039). **F.** Power spectrum of the contraTMS minus ipsiTMS time-series. Oscillatory peak was at 9 Hz and showed significant power (p < 0.0005, FDR-corrected: p = 0.0005). Note that 9 Hz oscillatory power was 71% higher in amplitude compared to the 10 Hz oscillation in the contraTMS condition, indicating an anti-phasic relationship.

To test if the oscillation in the ipsiTMS time-series is caused by a non-specific effect of TMS on both hemispheres, we tested whether oscillations in contra and ipsilateral TOJ time-series were consistent in phase. We therefore subtracted ipsiTMS time-series from the contraTMS time-series for each subject individually. Should both fluctuations in the time-series be caused by the same non-specific effect of TMS they should cancel out and the resulting contraTMS minus ipsiTMS time-series should show reduced amplitude. On the contrary, the resulting difference wave showed markedly higher fluctuations (71% increase in range) compared to the contraTMS time-series. By performing another frequency analysis, this time on the contraTMS minus ipsiTMS time-series, we could attribute this increase to a strong oscillatory component in the 9 Hz range (Figure 3F, 9 Hz: p < 0.00005, FDR corrected). The increase in oscillatory power in the difference wave is likely a result of subtracting two oscillations that share a common (or neighboring) frequency but are in anti-phasic relationship.

### 3. 10 Hz Phase analysis

To test this hypothesis we analyzed the 9 Hz phase angles of the ipsiTMS and contraTMS time-series (Figure 4A) as well as the difference wave across subjects. The frequency of 9 Hz was chosen because it lies right within the frequency ranges of the contra and ipsilateral peak components and both time series show highly significant power at this frequency (contraTMS: p = 0.00017; ipsiTMS: p = 0.0039). Additionally we performed a pairwise subtraction of the individual contraTMS and ipsiTMS phase angles (phase domain difference) to test if a possible anti-phasic relationship is visible on the single subject level. The complex FFT coefficients at 9 Hz were extracted to calculate individual phase angles (Figure 4B). We then compared these angles using Rayleigh’s test for non-uniformity testing the null hypothesis that the phase angles are randomly distributed. Phase angles were significantly clustered in ipsilateral as well as contralateral TOJ time-series (Figure 4B, contralateral: p = 0.000062, ipsilateral: p = 0.0017). We were further interested in the phase relationship between 9 Hz oscillations in contraTMS and ipsiTMS time-series (time domain difference). To test the two sets for phase opposition we subtracted the phase angles for each subject and performed Rayleigh’s test for non-uniformity on the resulting phase differences. Phase difference angles were significantly clustered at 180 degrees, indicating anti-phasic oscillations at opposing hemispheres (CI lower bound: 120.5 degrees, upper bound: 199.8 degrees, mean: 160.2 degrees, p < 0.01). The difference wave was analyzed in an identical fashion as contraTMS and ipsiTMS time-series, showing highly significant 9 Hz phase clustering (p=0.000001).

**Figure 4.**
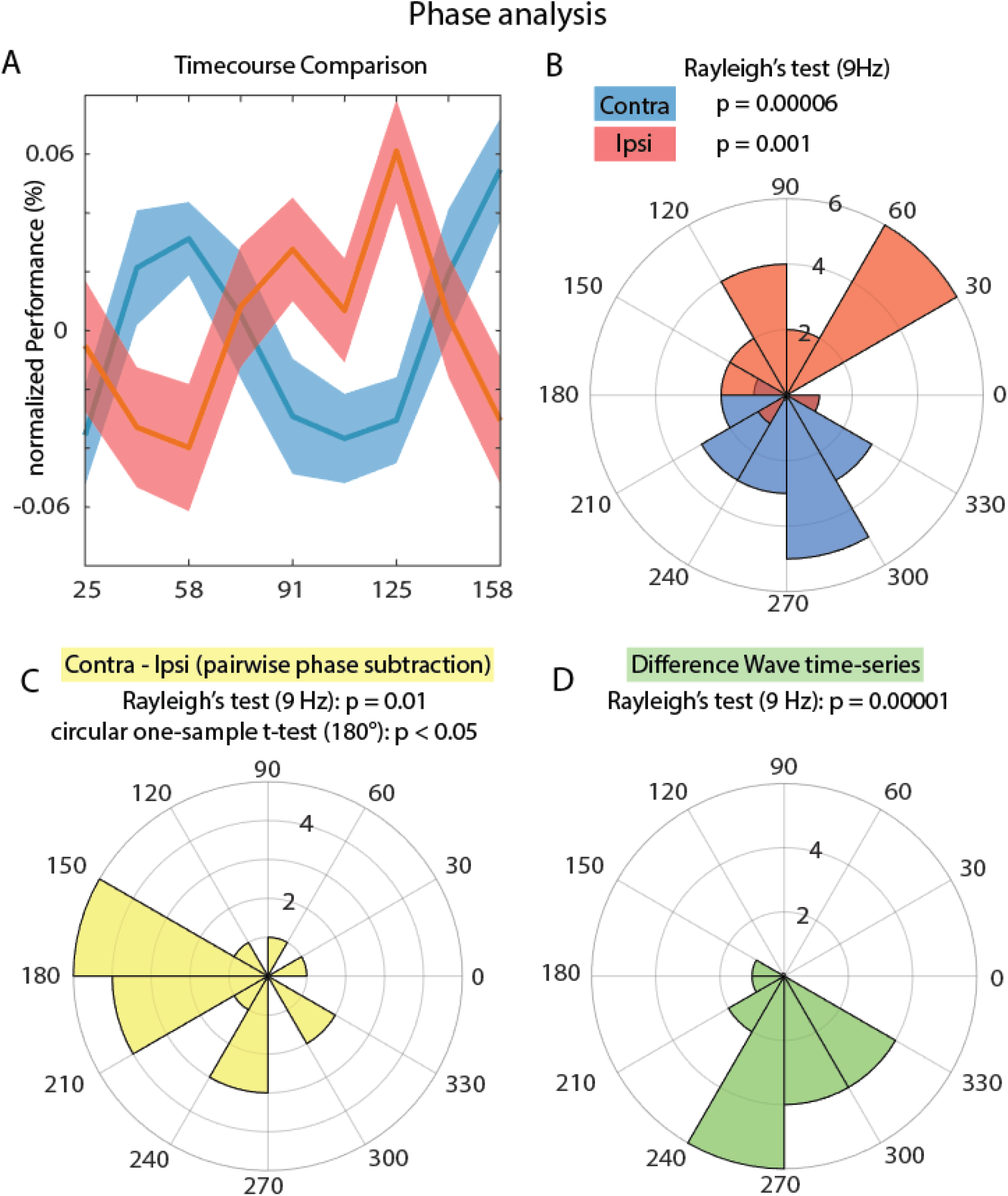
9 Hz Phase Analysis. **A.** Time Course of normalized contraTMS (stimulated hemifield) and ipsiTMS (non-stimulated hemifield) time courses for direct comparison. Note the clear antiphasic relationship between the two. **B.** 9 Hz phase angles of contraTMS (blue) and ipsiTMS (red) time-series. Rayleigh’s test reveals significant phase clustering for both conditions. **C.** Pairwise subtraction of 9 Hz contraTMS minus ipsiTMS phase angles (phase domain difference). Phase angles were significantly clustered around 180 degrees indicating an antiphasic relationship between individual contraTMS and ipsiTMS time-series. **D**. 9 Hz Phase analysis of the difference wave (contraTMS minus ipsiTMS, time domain difference). Phase angles were significantly clustered.

## Discussion

We tested the influence of α-TMS on subsequent temporal order judgements at varying SOA’s. 5 Pulses at 10 Hz were administered over left or right occipital cortex. α-TMS was intended to entrain α-oscillations in a local neural population. We probed temporal order perception of two Gabor patches, at the spatial location presumably affected by the TMS entrainment, at 9 SOA’s between 25 ms and 158 ms after the last TMS-pulse. Behavioral performance at every SOA was averaged to obtain a 133 ms long TOJ time-series. The frequency analysis of the TOJ time-series contralateral to TMS revealed a strong oscillation at our entrainment frequency of 10 Hz. We found no evidence of potentially confounding effects of TMS on stimulus visibility in our control conditions. In line with previous accounts of discrete perception we hypothesize that the rhythmic modulation in TOJ was caused by a TMS-evoked entrainment of occipital α-oscillations. The phase of α-oscillations has previously been related to so-called “perceptual windows” that serve to discretize visual input into compressed packages. Here we specifically tested the hypothesis that this compression leads to a deterioration of temporal order information. Depending on the relative timing to the last TMS pulse (SOA) the two stimuli fall either in the same or separate perceptual windows, leading to decreased or enhanced TOJ performance respectively. Our findings are in line with a “hard” theory of discrete perception suggesting that temporal order information is limited within perceptual windows. We provide causal evidence for an involvement of occipital alpha oscillations in this process.

The idea of discrete sampling in the visual system has received growing interest in the last years. Correlational evidence for an involvement of the occipital α-rhythm in the discretization of visual input is frequent (Busch et al., 2009; Haegens et al., 2011; Lőrincz et al., 2009; Milton & Pleydell-Pearce, 2016; Samaha & Postle, 2015; Valera et al., 1981; VanRullen, 2016; VanRullen & Koch, 2003; Vijayan & Kopell, 2012). Yet it was not clear if the α-rhythm merely modulates excitability, leading to continuous fluctuations in i.e. visual performance or if it implements discrete non-overlapping perceptual windows. These two notions can be thought of as “soft” and “hard” versions of the discrete perception idea (VanRullen, 2016). Evidence for the latter exists but is rarely of causal nature (Chakravarthi & Vanrullen, 2012; Milton & Pleydell-Pearce, 2016; Morand et al., 2015; Samaha & Postle, 2015; Valera et al., 1981; Wutz et al., 2014). Recent work shows that the α-rhythm can be causally modulated via visual entrainment, leading to fluctuations in temporal parsing performance (Ronconi et al., 2018; Chota & VanRullen, 2019). While these studies help to link α-oscillations and perception, they are limited since a visual entrainer passes various processing stages e.g. the LGN before arriving at the primary visual cortex. The LGN is hypothesized to project not only to V1, but also directly to higher cortical areas like V2 and V3 (Schmid et al., 2010). Strictly speaking every possible target of visual entrainment could serve as a potential source for the behavioral observations previously reported (Mathewson et al., 2010; Ronconi et al., 2018; Spaak et al., 2014; Chota & VanRullen, 2019). TMS allows for a direct and locally confined interaction with endogenous cortical rhythms (Romei et al., 2010; Thut et al., 2011). As the target of our entrainment is confined to the early occipital cortex we can say with relative certainty that the occipital α-rhythm gives rise to the perceptual effects demonstrated in this study. We therefore provide a direct link between occipital α-oscillations and temporal order judgements.

Former experiments have mostly used integration versus segregation tasks (IvS), such as the two-flash fusion paradigm, to quantify temporal parsing performance (Ronconi et al., 2018; Samaha & Postle, 2015; Valera et al., 1981). These tasks present two stimuli in quick succession and subsequently probe participants temporal segregation abilities (simultaneous vs. non-simultaneous, one vs. two). Perceptually it is principally sufficient to detect changes in luminance over time, irrespective of the stimulus characteristics. Flicker fusion experiments show that temporal changes can be detected at far higher frequencies than 10 Hz (Simonson & Brozek, 1952). TOJ’s however require precise perception of temporal relationships between individual stimuli and cannot be solved purely by identifying transient changes in luminance. Our findings suggest that fine grained temporal changes are detected at earlier stages of visual processing (e.g. LGN), whereas in early visual cortex (our target of entrainment) temporal order is resolved.

Another complicating factor of IvS tasks is that the oscillatory phase facilitatates or inhibits stimulus detection and ERP latencies, potentially biasing responses by hiding stimuli from perception or delaying their entry into consciousness (Busch et al., 2009; Fellinger et al., 2011). This raises the possibility that effects of phase on IvS performance are caused by inhibitory effects on single stimuli rather than time distortion effects caused by discrete sampling. One could also forward this objection against our paradigm.We therefore implemented several control conditions into our task. We hypothesized that if our findings were the result of phase-dependent α-inhibition, this should lead to a general reduction in the visibility of the target stimulus and reduce performance especially for low contrast conditions. Furthermore, such an effect would likely result in an increase in single stimulus reports for stimuli contralateral to TMS. Our control conditions however demonstrate that performance was comparable between TMS and Training conditions as well as contra and ipsilateral hemispheres in all contrast conditions, indicating that temporal order judgments could be performed under conditions of reduced visibility even under the effect of TMS. Even if one of the stimuli was occasionally hidden from perception entirely, we would remove them from the main TOJ-analysis as single stimulus reports. From our control results we conclude that modulations of stimulus visibility were not the main determinant of TOJ performance fluctuations.

While the effect of α-TMS on the contralateral visual field was somewhat expected we were surprised to find an oscillatory pattern also in the ipsilateral visual field. The ipsiTMS time-series showed a relatively weaker amplitude, oscillated at around 8 Hz and seemed to fluctuate in antiphase compared to its contralateral counterpart for at least 160 ms. We consider it unlikely that the TMS-pulse directly interacted with the contralateral hemisphere because first, all subjects reported phosphenes only in the contralateral visual field, and second, TMS pulses seemed to have opposing effects depending on the stimulus location. Previous work has shown that α-TMS can affect target detection performance in the visual field contra- and ipsilateral to the entrainment site, possibly explained by a transcallosal “push-pull” effect (Romei et al., 2010). Our findings suggest that α-phase is modulated by this network effect, potentially releasing one hemisphere from inhibition when the other hemisphere enters a state of inhibition. One possible interpretation of the frequency difference is that the ipsilateral oscillation reflects a slower attentional rhythm that is usually placed in the 3 to 8 Hz range (Fiebelkorn et al., 2018; Fiebelkorn & Kastner, 2019; VanRullen, 2013; VanRullen et al., 2007). Another possibility is that the change in frequency relates to differences in α-power between hemispheres. As mentioned before a push-pull effect might lead to a significant reduction in 10 Hz power in the ipsilateral hemisphere (Romei et al., 2010). Since this oscillatory frequency is likely relevant for stimulus processing the brain might try to compensate by modulating excitability at a lower frequency. Of course this is highly hypothetical and warrants further investigation.

The phase of α-oscillations is predictive of cortical excitability (Busch et al., 2009; Dugué et al., 2011), of neuronal firing rates (Haegens et al., 2011; Lőrincz et al., 2009; Vijayan & Kopell, 2012) and of the amplitude of gamma oscillations (Osipova et al., 2008; Voytek et al., 2010). As these neural signatures have been implicated in neuronal processing it seems logical that visual processing is concentrated on specific reoccurring intervals. The brain might use these naturally occurring periodicities, in the form of oscillations, to reduce the complexity of incoming information by compressing it into discrete packages. Our findings suggest that this compression results in the loss of temporal relationship between two stimuli. Note that the visual system is very robust to subsampling in the 100 ms range (VanRullen, Zoefel, & Ilhan, 2014). Presumably because visual information is highly redundant, allowing the visual system to reduce complexity via periodic discretization without losing too much relevant information.

In this study we successfully demonstrated that TMS entrainment at 10 Hz leads to a causal rhythmic modulation of temporal order judgements. This modulation was evident in the majority of subjects, shown by a strong inter-individual phase coherence in individual TOJ time-series. Furthermore we found that ipsilateral TOJ time-series were modulated in antiphase to their contralateral counterparts; this cross-hemispheric effect is unlikely to be caused directly by the TMS, but may result from a secondary trans-callosal pathway. We hypothesize that the visual system periodically discards temporal order information in order to reduce the complexity of incoming visual information. Further, we hypothesize that this mechanism is implemented by occipital oscillations in the α-range.

